# Changes in partner traits drive variation in plant–nectar robber interactions across habitats

**DOI:** 10.1101/2020.01.09.898692

**Authors:** Gordon Fitch, John Vandermeer

## Abstract

The frequency and outcome of biotic interactions commonly vary with environmental conditions, even without changes to community composition. Yet the drivers of such environmentally-mediated change in biotic interactions are poorly understood, limiting our ability to predict how environmental change will impact communities. Studying nectar robbery by stingless bees of *Odontonema cuspidatum* (Acanthaceae) in a coffee agroecosystem, we documented a temporally consistent difference in nectar robbing intensity between anthropogenic and seminatural habitats. Plants growing in coffee fields (anthropogenic habitat) experienced significantly more nectar robbery than plants growing in forest fragments (seminatural habitat). Using a combination of field surveys and manipulative experiments, we found that nectar robbery was higher in coffee fields primarily due to environmental effects on a) neighborhood floral context and b) *O. cuspidatum* floral traits. This led to both preferential foraging by nectar robbers in coffee fields, and to changes in foraging behavior on *O. cuspidatum* that increased robbery. Nectar robbery significantly reduced fruit set in *O. cuspidatum*. These results suggest that the effects of anthropogenic environmental change on species traits may be more important than its effect on species density in determining how interaction frequency and outcome are affected by such environmental change.

## Introduction

Understanding the effects of anthropogenic environmental change on biotic communities and ecosystem function is a key challenge for ecologists. Anthropogenic environmental change is resulting in striking biodiversity loss at a global scale (Dirzo et al., 2014; Matzke et al., 2011), with consequences for ecosystem function and ecosystem service provisioning at smaller scales (Hooper et al., 2012). But while the bulk of research on the effects of anthropogenic environmental change has focused on changes to community composition (Tylianakis et al., 2008; Valiente-Banuet et al., 2015), there is growing appreciation that the structure of interaction networks within communities can be altered by environmental change even when community composition is unaffected (Fig. 1; Poisot et al., 2015, 2017; Tylianakis et al., 2008; Tylianakis & Morris, 2017; Valiente-Banuet et al., 2015). Such interaction ‘rewiring’ may be mediated by changes to either the density (Fig. 1, Pathways A-B) or traits (Fig. 1, Pathways C-D) of one or more interacting species. These effects, moreover, may derive directly from altered environmental conditions (Fig. 1, Pathways A, D), or be mediated by environmentally-driven changes to community context (Fig. 1, Pathways B-C).

**Fig. 1.**
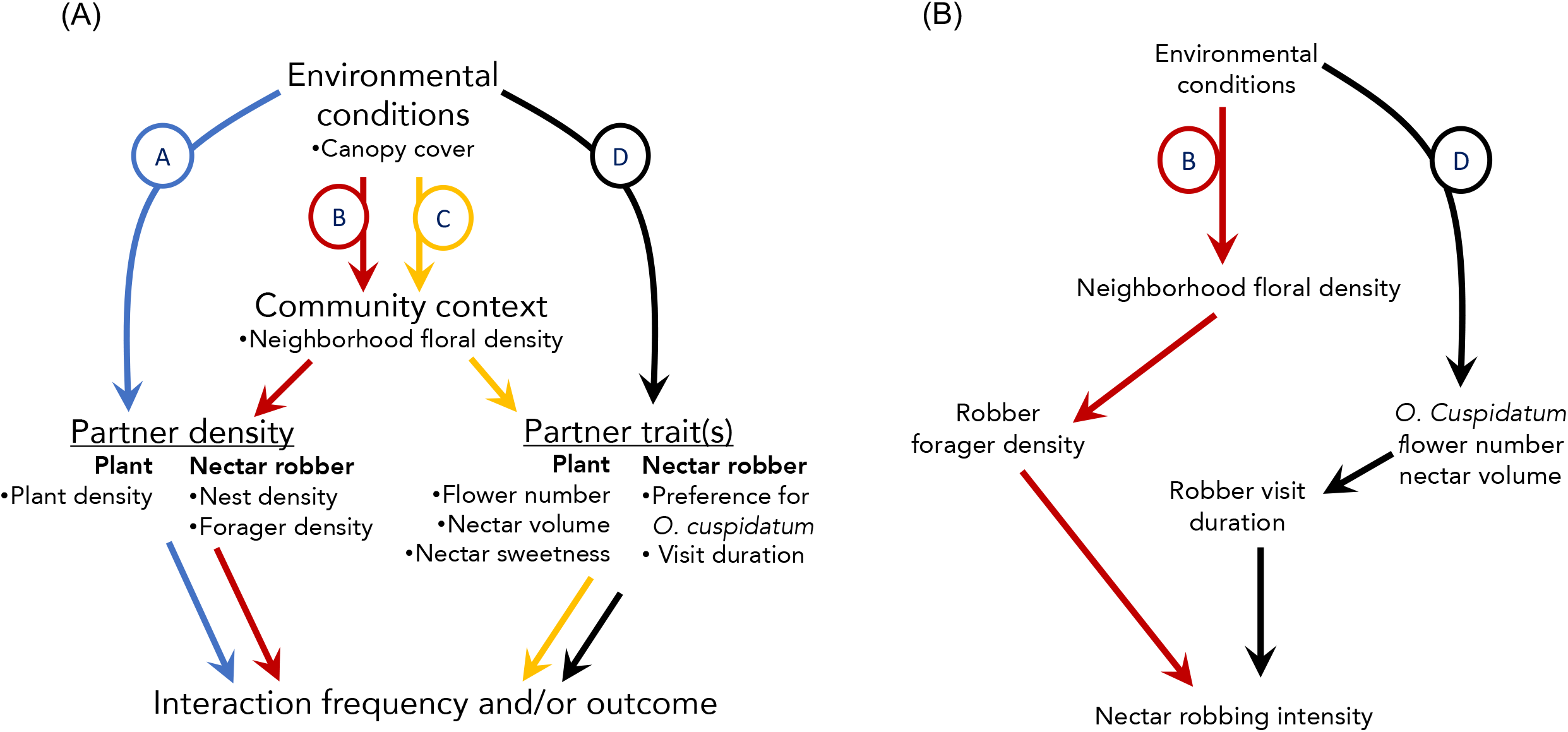
Pathways by which environmental change can result in changes to interactions without changing species composition. (A) Conceptual framework. Pathways are not mutually exclusive. Each line type represents one pathway. Bullet points indicate the aspects of each category assessed in this study. Pathways A-B: environmental conditions (e.g. insolation) directly affect the abundance (Pathway A) or one or more traits (Pathway B) of one or both of the interacting species. Pathways C-D: environmental conditions affect the traits or density of other species in the community, which in turn influences the abundance (Pathway C) or trait(s) (Pathway D) of one or more interacting species. (B) Schematic summary of results from this study, showing only pathways supported by the data.

Given the multiple pathways by which environmental change can impact interactions, it is perhaps not surprising that both theoretical (Valiente-Banuet et al., 2015) and empirical (Tylianakis et al., 2008; Valiente-Banuet et al., 2015) work suggest that changes in interaction structure are likely to occur more commonly, and more quickly, than species loss in response to environmental change. Yet, even where changes to the structure of interactions have been documented, we frequently lack understanding of the underlying drivers of these changes [Tylianakis et al. (2008), though see Fagundes et al. (2020) for an exception]. This limits our ability to predict how future environmental change will impact communities and ecosystems.

Nectar robbery (NR), in which a flower visitor extracts nectar from a flower via an opening other than the corolla mouth, is one interaction type that is likely to be strongly impacted by environmental context (Cuevas & Rosas-Guerrero, 2016; Irwin & Maloof, 2002; Morris, 1996), since it is generally a facultative behavior (Irwin et al., 2010; Morris, 1996; Richardson & Bronstein, 2012). The effects of NR on reproductive success of the robbed plant are variable; though negative, neutral, and positive effects have been reported (Burkle et al., 2007; Maloof & Inouye, 2000), negative effects, at least on components of female fitness, are most common (Irwin et al., 2001). The fitness outcome of NR for the robbed plant depends on the interaction of a number of factors, including plant mating system, the identity and foraging behavior of both robbers and legitimate pollinators, and the environmental (particularly floral) context in which the interaction occurs (Burkle et al., 2007; Maloof & Inouye, 2000; Morris, 1996).

Nectar-robbing intensity (NRI) – measured as the proportion of flowers that experience robbery – commonly varies both spatially (Cuevas & Rosas-Guerrero, 2016; Irwin & Maloof, 2002; Morris, 1996) and temporally (Cuevas & Rosas-Guerrero, 2016; Irwin & Maloof, 2002; Navarro, 2000). Multiple drivers of such variability have been postulated; these drivers are not mutually exclusive, and in some cases multiple drivers may be operating in tandem. Putative drivers include both direct responses of robber or plant to environmental conditions and responses mediated by the broader community. Direct responses may include variation in the density of robbers (Irwin & Maloof, 2002; Navarro, 2000) or the density, flower number, or nectar quality or quantity of the focal plant (Krupnick et al., 1999) due to environmental conditions. Community-mediated responses may include altered foraging behavior depending on the availability of alternative floral resources (Irwin et al., 2010; Irwin & Maloof, 2002) or density of other flower visitors (Roubik, 1982). Yet to date, there has been little work documenting which of these mechanisms operate in specific instances to generate variation in NRI. Without a mechanistic understanding of the ecological drivers of NR, it is difficult to predict the circumstances under which NR will occur and how it will be altered by environmental change.

In this study, we first assessed the intensity of NR by stingless bees of the shrub *Odontonema cuspidatum* (Nees) Kuntze (Acanthaceae) in a semi-natural habitat (forest fragments) and an anthropogenic one (coffee farm). We then evaluated the role of potential drivers of NR (Fig. 1) in generating habitat-based spatial heterogeneity in NRI. We also evaluated whether NR influenced either the likelihood of individual flowers setting fruit or plant-level reproductive output across both habitats. Our aim was to understand the extent to which variation in the intensity and outcome of NR across habitats is driven by 1) changes in population density of robber or plant (Fig. 1 Pathways A-B), 2) direct effects of environmental conditions (i.e. light availability) on one or more traits of either partner (Fig. 1 Pathway D), or 3) indirect effects on robber or plant traits via changes in community context (Fig. 1 Pathway C).

We interpret the term ‘trait’ to include both physical characteristics and behaviors, consistent with the definition used in the literature on trait-mediated indirect effects (e.g. Werner & Peacor 2003, Schmitz et al. 2004, Utsumi et al. 2010). Specifically, the traits we focus on are, for the plant, flower number and floral nectar characteristics and, for the nectar robbers, foraging behavior.

## Materials and methods

### Study system

This research was conducted at Finca Irlanda, a shaded, organic coffee farm, approximately 300 ha in size, located in the Soconusco region of southeastern Chiapas, Mexico. The farm ranges from 900-1150 masl; above ∼1000 masl it is comprised primarily of *Coffea arabica* plantations, with three small (<30 ha) forest fragments embedded within the farm. Forest fragments are characterized by a higher density and diversity of canopy trees in comparison to the coffee fields. As a result of the higher density of canopy trees in forest fragments, the amount of light reaching the ground is generally lower in the forest than the coffee farm (see Results). This difference in canopy cover represents a key environmental difference between these habitats.

Within this landscape, *O. cuspidatum*, a perennial shrub native to the region, grows both in areas under coffee cultivation and in forest fragments. In the study area, *O. cuspidatum* blooms primarily from June to August, in the early part of the rainy season. Slender red flowers, 2-2.5 cm long, are borne on indeterminate branching racemes; individual plants produce from 1 to approx. 30 racemes, and each raceme holds from approx. 10 to hundreds of flowers (G. Fitch unpublished data). *Odontonema cuspidatum* is self-fertile but requires animal pollination for fertilization, due to spatial separation of anthers and stigma (G. Fitch unpublished data). Hummingbirds, particularly the blue-tailed hummingbird (*Amazilia cyanura*), are the most frequent legitimate floral visitors (G. Fitch unpublished data); this, together with the flower’s morphology, suggests that hummingbirds are the primary pollinators of *O. cuspidatum*. The flowers also attract a wide range of nectar-feeding insects, most of which engage exclusively in nectar robbery, extracting nectar from animal-made holes in the base of the corolla tube. Primary nectar robbers – i.e. those that make the hole themselves, hereafter ‘PNR’ – comprise two species of stingless bee in the genus *Trigona* (*T. fulviventris* and *T. nigerrima*; Hymenoptera: Apidae: Meliponini). Fertilized flowers produce explosively dehiscent capsules (Daniel, 1995).

### Data collection

#### Spatiotemporal patterns of nectar robbing intensity

Within a 25ha area that included both coffee fields and forest fragments, we haphazardly selected 109 individual *O. cuspidatum* for inclusion in the study. This represents ∼50% of all individuals found in the survey area. This 25ha section of the farm was selected because it included the principal forest fragments contained within the farm’s boundaries, as well as high densities of *O. cuspidatum*. Because of the spatial arrangement of the forest fragments, each plant was within 500 m of a habitat edge. All plants were individually labeled and followed through both years of the study, except that thirty-three plants surveyed in 2017 either died or did not flower in 2018, and an additional 15 plants that flowered in 2018 but not 2017 were monitored in 2018 only. We recorded the GPS coordinates of each plant. Distance between plants included in the study ranged from 10-2200 m.

In 2017-2018, plants were surveyed for NR weekly for the duration of the flowering period. At each survey, all inflorescences with open flowers were surveyed. Flowers >1.5 cm long were checked for evidence of NR (characteristic hole at corolla base), and the number of robbed and unrobbed flowers on each inflorescence was recorded. The 1.5 cm cutoff was chosen because flowers of this length were generally within 2 days of opening, and prior to this stage flowers experienced minimal NR (G. Fitch unpublished data). In 2018 only, each monitored inflorescence was individually tagged.

### Putative drivers of nectar robbing intensity

#### Odontonema cuspidatum density

To determine the density of *O. cuspidatum* in each habitat, in June 2018 we counted the number of *O. cuspidatum* inflorescences, at any stage of development, within a 20 m radius of each of our target plants.

#### Odontonema cuspidatum floral traits

Each time we surveyed plants for NR, we recorded the total number of flowers present on the plant, for our measure of per-observation flower number. At the end of the flowering period, the season-long total number of flowers produced was determined by counting all mature fruits and persistent ovaries (i.e. flowers that had not set fruit) on each plant.

On a subset of monitored plants (49 in 2017, 19 in 2018 with 7 included in both years), we assessed nectar volume and sugar content. Because standing nectar crop was minimal in unbagged flowers, to measure nectar content we covered 2 inflorescences/plant with a bag made of 0.5 cm tulle mesh to exclude floral visitors. During the flowering period, we checked bagged inflorescences for open flowers 2x/week. To assess nectar volume, we removed all nectar from a flower using a 75 μ L microcapillary tube (Drummond Scientific, Broomall PA), then measured the height of the nectar in the tube using digital calipers (Thomas Scientific, Swedesboro NJ) and converted this measure to nectar volume. We used a pocket refractometer (Eclipse 45-81, Bellingham & Stanley, Tunbridge Wells UK) to assess the nectar sugar content of each sample. From each plant, we assessed nectar characteristics of at least 4 flowers (range: 4-32 flowers, mean ± SE: 12.9 ± 0.3 flowers).

### Primary nectar robber (PNR) density and foraging behavior

To determine PNR population density in each habitat, in 2018 we conducted surveys for PNR nests. Surveys were conducted along 30 m x 10 m transects oriented in one of the cardinal directions and centered on a target *O. cuspidatum* plant. We conducted surveys along 32 transects, 16 in coffee and 16 in forest. Within a habitat, target plants for surveys were selected at random, except once a plant was selected, all other plants falling within the transect were excluded from selection, so no transects overlapped. Collectively, these transects encompassed 44 monitored *O. cuspidatum* plants. The PNR species nest either in trees or in the ground at the base of trees (Fierro et al., 2012), so our nest search focused on trees ≥15 cm dbh (Hubbell & Johnson, 1977). On all such trees within each transect, we scanned the trunk and major limbs from 0-20m above the ground for evidence of nesting. Surveys were conducted between 0700-1100, when nest activity was highest. Both species of PNR of *O. cuspidatum* in the study area have prominent nests with high activity levels, so we are reasonably confident that we located all nests of these species within our transects.

To determine PNR forager density, in addition to nest density, in 2017 and 2018 we surveyed local PNR abundance for each focal *O. cuspidatum* plant. Surveys occurred during peak bee activity (0700-1100 h) on sunny days. Each survey consisted of two 10-min periods. The first focused on flowers from 2-5 inflorescences of the focal *O. cuspidatum*, and the second focused on flowers within 10 m of the focal plant, with a 10-min break in between surveys. Prior to beginning the survey, we counted the number of open flowers on focal inflorescences, and located all bee-attractive flowering plants in the 10 m neighborhood. During surveys, all insect flower visitors were caught using resealable plastic bags and held, one insect per bag, for the duration of the survey. At the end of each 10-min survey, captured insects were identified to species or morphospecies and then released. Insects were classified as primary nectar robbers, secondary nectar robbers, legitimate visitors, or unknown, based on prior observation of insect visitation to *O. cuspidatum*. These observations provided us with several measures of PNR forager abundance: total site-level abundance, abundance on focal *O. cuspidatum*, per-flower abundance on focal *O. cuspidatum*, abundance on non-focal flowers, and the proportional abundance on target plants (number of individuals caught on target plants divided by total number of individuals caught during that survey). We interpret the latter metrics as a measure of PNR foraging behavior, while we consider the others to be measures of PNR foraging behavior.

In 2018, we assessed within-plant PNR foraging behavior on 23 monitored plants, 13 in coffee and 10 in forest. On each plant, we monitored five foraging bouts by individual PNRs at focal *O. cuspidatum* plants, recording the species of PNR, the length of time spent on the inflorescence, and the number of flowers robbed. We then calculated the proportion of potentially robbable flowers that had been robbed.

### Environmental conditions – Canopy cover

We measured canopy cover directly above the crown of each focal plant (N=109) in June 2017 (June 2018 for plants added in 2018) using CanopyApp 1.0.3 (University of New Hampshire, Durham, NH USA).

### Community context – Floral resource availability

In 2017, floral surveys were conducted to determine floral resource availability in the neighborhood of 95 focal plants. For each plant, four 10 m x 2 m transects were established, each one beginning at the focal plant and extending 10 m in one of the cardinal directions. Along each transect, all blooms were counted and identified to species or genus. Surveys occurred once for each focal plant, near the peak bloom for each focal plant. In some cases, transects for two plants partially overlapped; in these cases, bloom tallies for the overlapping portions of the transects were included in the total for both plants’ transect.

#### Effect of nectar robbery on reproduction

For monitored plants that had >20 flowers and had both robbed and unrobbed flowers (N=44), we evaluated the difference in probability of producing fruit and in seed set between robbed and unrobbed flowers. Flowers were marked with either red (robbed) or blue (unrobbed) nail polish on the pedicel. Robbed flowers were marked either on the day before they were to open or soon after opening, while unrobbed flowers were marked only as they were beginning to senesce, to ensure that robbing occurred before pollination and that flowers marked as unrobbed were not subsequently robbed. The number of marked flowers of each type (robbed and unrobbed) varied across plants depending on the availability of robbed and unrobbed flowers (robbed flowers: range = 1-26 flowers, mean ± SE = 4.7 ± 0.7 flowers; unrobbed flowers: range = 1-16 flowers, mean ± SE = 2.9 ± 0.4 flowers).

We assessed fruit set by counting the number of fruit and number of persistent ovaries on mature inflorescences. The fate of each marked robbed or unrobbed flower was recorded. Inflorescences that had been damaged by insect herbivores were excluded from further analyses. To measure seed set, up to 5 fruits (in 2017) or all undamaged fruits (in 2018) were collected from each inflorescence. These fruits, segregated by plant, were placed in small bags made of 0.5 cm mesh fabric and left in a drying oven at 50 °C until all had dehisced, at which time seeds were counted.

### Data analysis

All analyses were conducted in R v.3.5.1 (R Core Team, 2018). All models were checked for conformity to assumptions.

#### Influence of habitat on nectar robbing intensity

We evaluated the influence of habitat on NRI using GLMMs with habitat and year as fixed effects and plant and date as random effects. We separately considered three components of NRI: number of open flowers robbed, number of unopened flowers robbed, and total number of flowers robbed. In all cases, we included log(total number of flowers of that class) as an offset, and used a Poisson error distribution with log-link function. To check for spatial autocorrelation in NRI, we fit a parallel set of GLMs, and calculated Moran’s I for the residuals of each of these models using package ‘ape’ (Paradis & Schliep, 2018). We found no evidence for spatial autocorrelation in these data (p > 0.1 in all cases).

#### Influence of habitat on putative drivers of nectar robbing intensity

We tested for a significant effect of habitat on the following putative drivers of NRI: focal plant flower number, focal plant nectar volume and nectar sugar content, canopy cover, neighborhood floral density, PNR density at focal plants and in the 10-m neighborhood, and PNR nest density. We initially considered several floral neighborhood metrics: floral richness, total flower density, and *O. cuspidatum* flower density. These metrics were strongly correlated and total flower density was the best predictor of NRI, so for all analyses we used total floral density to assess effects of floral neighborhood. Focal plant flower number (assessed as the total number of flowers produced over the flowering period) and PNR nest density were fitted with a GLM with Poisson distribution and log-link function. Focal plant nectar volume and nectar sugar content, as well as canopy cover, were fitted using linear models. For other putative drivers, we had multiple measures per plant, and therefore used GLMMs with plant as a random effect and a Poisson distribution with log-link function. We calculated Moran’s I for the residuals of all models to check for spatial autocorrelation. For drivers with significant spatial autocorrelation, we used the ‘spdep’ package to calculate Moran eigenvectors (Bivand & Wong, 2018). These eigenvectors were included as additional predictors in an updated model. Where significant spatial autocorrelation was found, the estimates we provide for the effect of habitat on the relevant driver come from the model including Moran eigenvectors. To evaluate significant effects, all p-values were adjusted for multiple comparisons using the Bonferroni correction.

We also tested whether canopy cover (and therefore light availability) could account for differences in floral traits (flower number and flower nectar volume) between habitats. For flower number, we used a GLM with negative-binomial distribution, and for nectar volume we used a linear mixed-effects model with plant as a random effect.

#### Influence of putative drivers on nectar robbing intensity

The number of plant-year combinations for which we had observations of a particular putative driver varied substantially across drivers. This variation precluded analysis using structural equation modeling, because the combined dataset for all putative drivers contained too few observations for meaningful analysis given the number of drivers (Kline, 2015). Therefore, we initially assessed the relationship between NRI and each putative driver that differed significantly between habitats (that is, all the boldface variables listed in Table 1) separately, then included those drivers that had an effect on NRI at Bonferroni-corrected p ≤ 0.2 in a combined model (see below). All models used the number of robbed flowers as the response variable, and included log(total number of flowers surveyed) as an offset; error distributions were either Poisson or negative binomial, depending on whether the data were overdispersed, and used a log-link function.

**Table 1.**
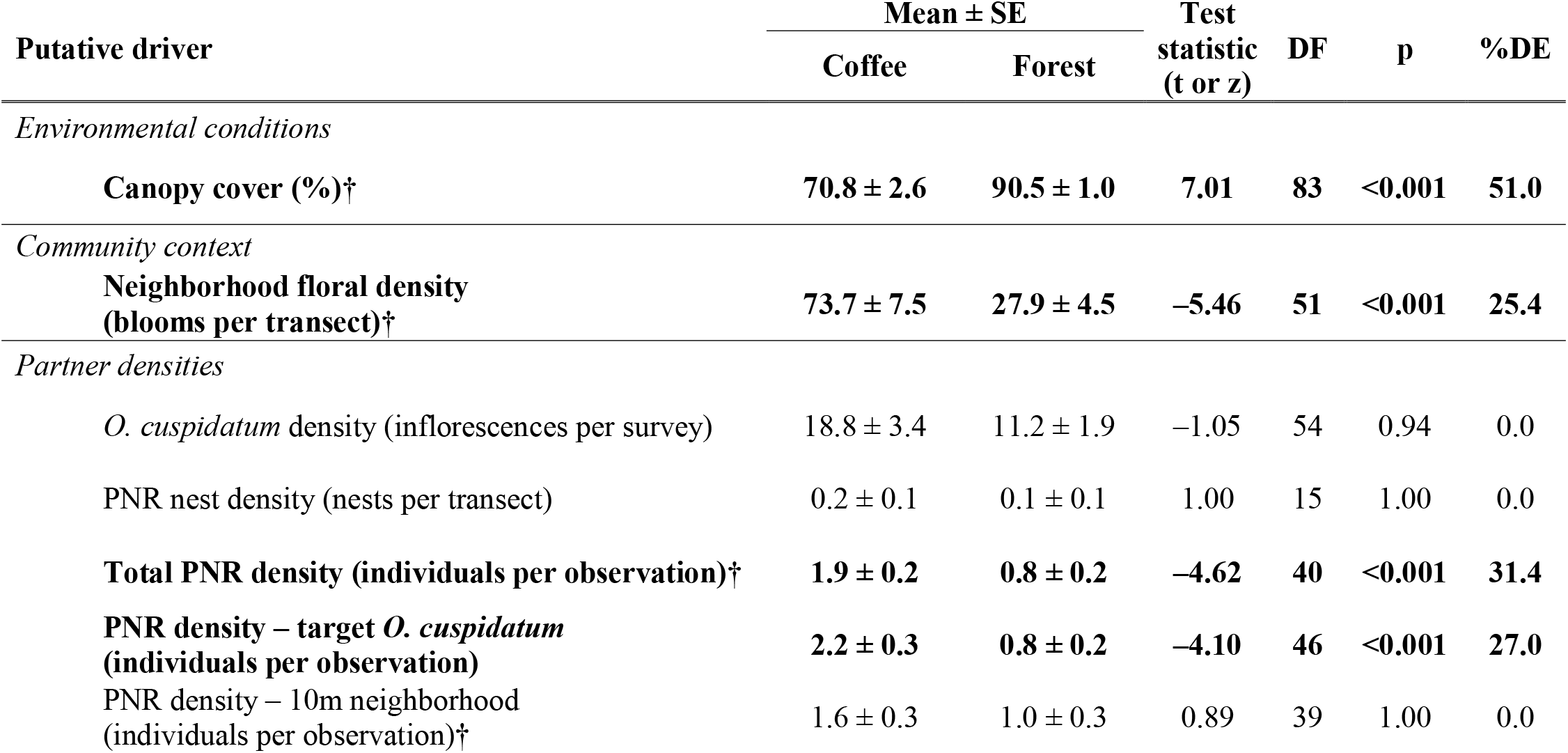
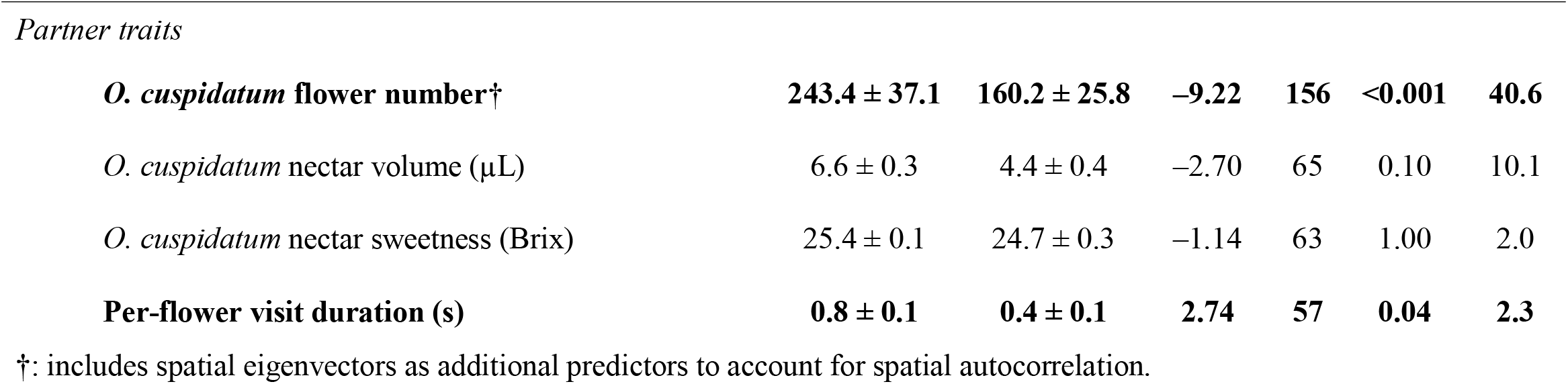
Relationship between habitat and putative drivers of nectar robbery. Model results are from generalized linear models (canopy cover, neighborhood floral density, *O. cuspidatum* density, and PNR nest density) or generalized linear mixed-effects models with plant as a random effect (all other variables). PNR: primary nectar robber; DF: residual degrees of freedom; %DE: percent of the null deviance in the relevant response variable explained by habitat; Boldface indicates a significant difference between habitats in levels the putative driver, at p < 0.05. All p-values are Bonferroni-corrected for multiple comparisons.

| Putative driver | Mean ± SE | | Test statistic (t or z) | DF | p | %DE |
| --- | --- | --- | --- | --- | --- | --- |
|  | Coffee | Forest |  |  |  |  |
| *Environmental conditions* |  |  |  |  |  |  |
| Canopy cover (%)† | 70.8 ± 2.6 | 90.5 ± 1.0 | 7.01 | 83 | <0.001 | 51.0 |
| *Community context* |  |  |  |  |  |  |
| Neighborhood floral density (blooms per transect)† | 73.7 ± 7.5 | 27.9 ± 4.5 | −5.46 | 51 | <0.001 | 25.4 |
| *Partner densities* |  |  |  |  |  |  |
| *O. cuspidatum* density (inflorescences per survey) | 18.8 ± 3.4 | 11.2 ± 1.9 | −1.05 | 54 | 0.94 | 0.0 |
| PNR nest density (nests per transect) | 0.2 ± 0.1 | 0.1 ± 0.1 | 1.00 | 15 | 1.00 | 0.0 |
| Total PNR density (individuals per observation)† | 1.9 ± 0.2 | 0.8 ± 0.2 | −4.62 | 40 | <0.001 | 31.4 |
| PNR density – target *O. cuspidatum* (individuals per observation) | 2.2 ± 0.3 | 0.8 ± 0.2 | −4.10 | 46 | <0.001 | 27.0 |
| PNR density – 10m neighborhood (individuals per observation)† | 1.6 ± 0.3 | 1.0 ± 0.3 | 0.89 | 39 | 1.00 | 0.0 |
| *Partner traits* |  |  |  |  |  |  |
| ***O. cuspidatum* flower number†** | **243.4 ± 37.1** | **160.2 ± 25.8** | **−9.22** | **156** | **<0.001** | **40.6** |
| *O. cuspidatum* nectar volume (μL) | 6.6 ± 0.3 | 4.4 ± 0.4 | −2.70 | 65 | 0.10 | 10.1 |
| *O. cuspidatum* nectar sweetness (Brix) | 25.4 ± 0.1 | 24.7 ± 0.3 | −1.14 | 63 | 1.00 | 2.0 |
| **Per-flower visit duration (s)** | **0.8 ± 0.1** | **0.4 ± 0.1** | **2.74** | **57** | **0.04** | **2.3** |

†: includes spatial eigenvectors as additional predictors to account for spatial autocorrelation.

The relationship between flower visitation and both flower number and floral neighborhood is often nonlinear and unimodal (Ghazoul, 2006; Rathcke, 1983). Beyond unimodality, we did not have *a priori* expectations for the relationship between these drivers and NR. Therefore, to assess their effect on NRI, we used general additive models (GAMs). To test for nonlinearity, we compared model versions with linear and smoothed relationships using AICc. In both cases, the nonlinear model indicated a unimodal relationship and improved fit over the linear model (for flower number, ΔAICc = 19; for floral neighborhood ΔAICc = 25). For all other drivers we assumed a linear response of NR; we used a GLM to evaluate the effect of canopy cover and floral neighborhood and GLMMs to evaluate the effect of all other drivers (since for these drivers we had multiple observations per plant).

These single-predictor models indicated that at the p ≤ 0.2 level, the putative drivers that affected NRI were focal plant flower number, neighborhood floral density, and robber density at focal plants. To test whether these putative drivers had independent effects, we used a GAM with all three predictors; smooth terms were applied to flower number and floral neighborhood, while robber density was constrained to a linear function.

Based on GAM results for the relationship between NRI and neighborhood floral density, we classified neighborhood floral densities as low (less than the lower bound of the 95% confidence interval for the predicted maximum of the function relating NRI to floral density), moderate (within the 95% confidence interval), or high (higher than the upper bound of the 95% confidence interval).

#### Effects of nectar robbery and habitat on Odontonema cuspidatum reproduction

To determine the effect of NR on *O. cuspidatum* reproduction, we compared differences in fruit set between robbed and unrobbed flowers using a binomial GLMM with plant as a random effect. We additionally tested for an effect of habitat on fruit set, and of differences in the effect of NR on fruit set between habitats, by including habitat and a robbed status × habitat interaction term as fixed effects. To control for differences across plants in their ability to produce fruit, independent of the effects of NR, we only included plants for which we had data on the fate of both robbed and unrobbed flowers in a single year (N = 44).

To test the effect of NRI and habitat on measures of reproductive output, we examined fruit set, seed set, and seeds produced per plant. To model fruit set, we used a Poisson generalized linear mixed-effects model (GLMM) with number of fruits as the response variable, offset by log(total number of flowers produced). Fixed effects were habitat, NRI, and a habitat × NRI interaction term; plant was included as a random effect. Seed set and seeds per plant were assessed only in 2018, so for those metrics we used a negative binomial generalized linear model (GLM) with number of seeds as the response; the model for seed set additionally included log(total number of flowers) as an offset. Because there was no effect of the habitat × NRI interaction term in the model for fruit set, this predictor was omitted from our models of seed set and seeds per plant.

## Results

### Spatiotemporal patterns in nectar robbing intensity (NRI)

Plants growing in forest fragments were robbed significantly less than those growing in coffee, consistent across years (proportion of flowers robbed in forest: 0.29 ± 0.02; in coffee: 0.42 ± 0.02; z = –2.26, p = 0.02). Across both habitats, NRI was significantly higher in 2018 than 2017 (2017: 0.32 ± 0.02; 2018: 0.46 ± 0.01; z = 8.70, p < 0.001) and, in both years, increased over the survey period (ß = 0.19±0.02, z = 10.49, p < 0.001).

### Relationships between putative drivers and habitat

#### Environmental conditions and community context

Canopy cover over target *O. cuspidatum* was significantly higher in forest fragments than coffee (Table 1). Neighborhood floral density was significantly higher in coffee than in forest fragments (Table 1).

#### Partner densities

Density of *O. cuspidatum* did not differ between habitats (Table 1). Density of primary nectar robbers (PNRs), on the other hand, showed a complex response to habitat. There was no difference in nest density of the two PNR species between habitats (Table 1). Nevertheless, PNR density at monitored *O. cuspidatum* plants was higher in coffee than in forest (Table 1), suggesting greater forager density in coffee. However, PNR density at all other flowers within 10 m of focal plants (most of which received legitimate visits from PNRs) was equivalent across habitats (Table 1). Because of the high density of PNRs at target *O. cuspidatum*, total PNR density was still significantly higher in coffee than in forest (Table 1).

#### Partner traits

*Odontonema cuspidatum* flower number was significantly lower in forest (Table 1). Nectar volume and sweetness were both lower for plants growing in coffee, but these differences were not significant (Table 1). These changes in floral traits stem, at least in part, from reduced light availability in forest fragments, as canopy cover (significantly higher in forest fragments) had a negative effect on both total flower number (ß = –0.31±0.08, z = –3.72, p < 0.001) and nectar volume (ß = –0.19±0.08, t = –2.82, p = 0.03).

Primary nectar robber foraging behavior at *O. cuspidatum* plants differed between habitats. On a per-flower basis, PNRs spent more time per foraging bout on plants growing in coffee (Table 1).

### Relationships between putative drivers and nectar robbery

Nectar robbery responded nonlinearly to both *O. cuspidatum* flower number and neighborhood floral density (Fig. 2, Table 2). In both cases, NRI had a unimodal response, initially increasing, then decreasing as floral availability increased further (Fig. 2). For *O. cuspidatum* flower number, the position of the predicted maximum was 645 blooms [95% confidence interval (CI): 467-850 blooms; Fig. 2A]. This is larger than the mean flower number for individuals from either forest fragments (160 ± 26) or coffee fields (243 ± 37), indicating that for most of the surveyed plants, increasing flower number leads to increased NRI.

**Fig. 2.**
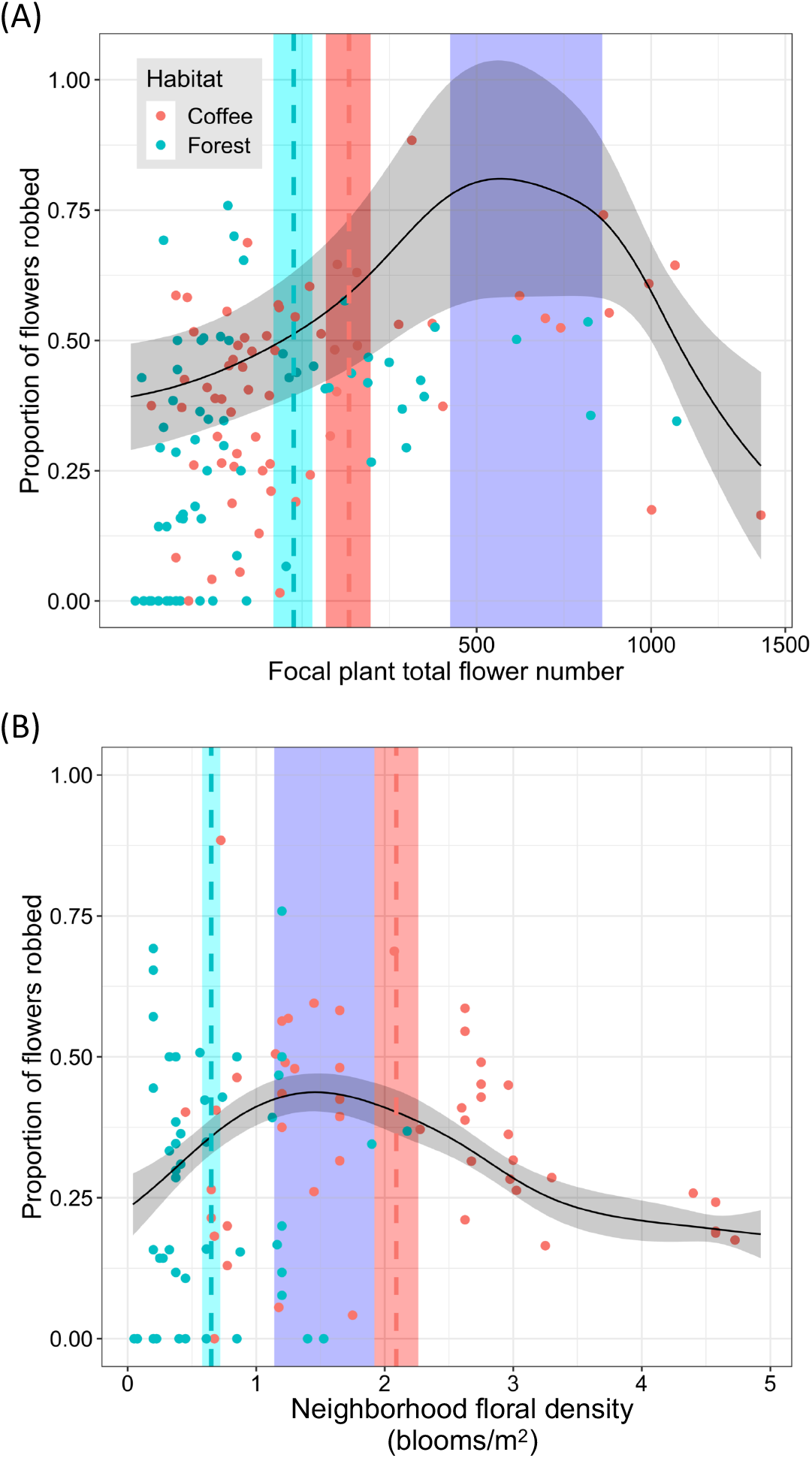
Relationship between nectar robbing intensity (NRI) of *O. cuspidatum* and (A) focal plant flower number and (B) floral neighborhood. In both panels, points represent individual plants, colored according to habitat. Black line represents best-fit from a general additive model; gray-shaded area represents standard error. Blue shaded area represents 95% confidence interval about (A) *O. cuspidatum* flower number or (B) the neighborhood floral density where maximum NRI is predicted. Dashed vertical lines and shaded area represent the mean ± standard error for (A) focal plant flower number and (B) neighborhood floral density in each habitat. ‘Neighborhood floral density’ refers to the mean number of blooms of all species per transect. In (A), the x-axis is square-root transformed to improve legibility.

**Table 2.**
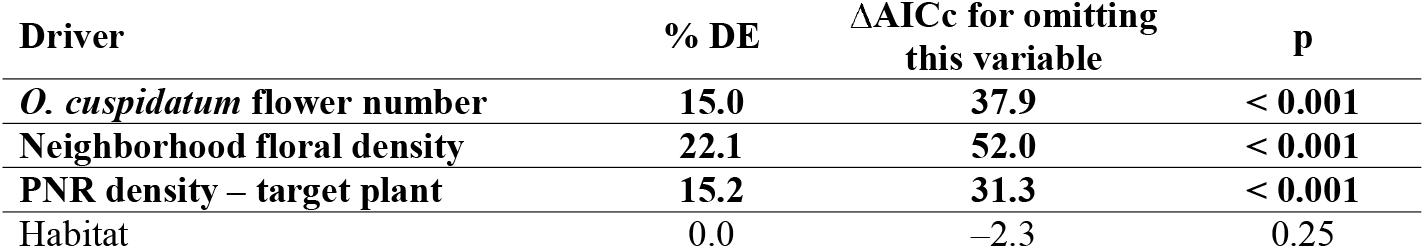
Relative importance of the significant drivers of nectar robbing intensity (NRI). Results from a general additive model combining all of the putative drivers that significantly influenced NRI when considered independently. Boldface indicates a significant effect on NRI at p < 0.05. % DE: percent of the null deviance in NRI explained by each driver; PNR: primary nectar robber

For neighborhood floral density, the predicted maximum was 1.8 blooms/m^2^ (95% CI: 1.1–1.9 blooms/m^2^; Fig. 2B). Significantly, the neighborhoods of most forest-growing *O. cuspidatum* had low floral densities (i.e. below the lower bound of the 95% CI for the predicted maximum: 38 plants below, 11 within, 3 above). The floral neighborhoods of coffee-growing plants varied more in density (10 plants below the CI; 16 within; 23 above; Fig. 2B), with a substantially higher percentage of coffee-growing plants found in floral neighborhoods within the 95% CI for peak NRI (33% versus 22%). Most (65%) of the plants whose neighborhood floral density fell within the 95% CI for predicted maximal NRI grew in coffee.

The other putative driver that significantly affected NRI was PNR density on target plants, which was higher for coffee-growing plants (Table 1). When these three significant drivers were combined into a single model, all three retained their significance level and showed a qualitatively similar effect on NRI as when they were considered independently (Table 2). Moreover, when habitat was added as a linear predictor to this model, it did not have a significant effect on NRI and did not improve model fit (Table 2).

### Effects of nectar robbery and habitat on Odontonema cuspidatum reproduction

Nectar robbery significantly reduced the probability of a flower setting fruit, from 0.32 ± 0.04 (mean ± SE) for unrobbed flowers to 0.18 ± 0.03 for robbed flowers, representing a 43% decrease in fruit set (z = 3.41, p < 0.001). The effect of NR on probability of setting fruit was equivalent between habitats (robbed status × habitat interaction: ß = 0.16 ± 0.60, z = 0.27, p = 0.8), as was the overall probability of setting fruit (coffee: 0.23 ± 0.03; forest: 0.23 ± 0.04; z = – 0.10, p = 0.9).

Consistent with the negative effect of NR on the probability of individual flowers setting fruit, we found a significant negative relationship between NRI and fruit set at the plant level (Table 3). This effect was consistent across habitats, and fruit set did not differ between habitats (coffee: 0.12 ± 0.01; forest: 0.11 ± 0.01; Table 3). By contrast, neither seed set nor seeds per plant were influenced by NRI. Seed set was higher in forest-growing plants, though due to the smaller number of flowers produced by forest-growing plants, seeds per plant did not differ between habitats (Table 3).

**Table 3.** Relationship between reproductive output and nectar robbing and habitat. DF: residual degrees of freedom; %DE: percent of the null deviance in the measure of reproductive output that is explained by each predictor; NRI: nectar robbery intensity. Boldface indicates statistical significance at p < 0.05

| Predictor | Coefficient<br>($\beta \pm SE$) | Test statistic<br>(t or z) | DF | %DE |
| --- | --- | --- | --- | --- |
| Fruit set (GLMM) |  |  |  |  |
| **NRI** | **$-0.63 \pm 0.21^{***}$** | **-2.94** | **123** | **8.0** |
| Habitat (Forest) | $-0.34 \pm 0.23$ | -1.45 | 123 | 0.8 |
| NRI × Habitat | $0.31 \pm 0.31$ | 1.00 | 123 | 0.0 |
| Seed set (2018 only; GLM) |  |  |  |  |
| NRI | $-0.33 \pm 0.36$ | -0.91 | 56 | 1.2 |
| Habitat (Forest) | $0.38 \pm 0.20$ | 1.92 | 56 | 5.5 |
| Seeds per plant (2018 only; GLM) |  |  |  |  |
| NRI | $0.88 \pm 0.52$ | 1.71 | 56 | 2.7 |
| Habitat (Forest) | $-0.05 \pm 0.30$ | -0.17 | 56 | 1.5 |

. $p = 0.05$ , \*\*\* $p < 0.001$ .

## Discussion

Changes in the frequency or outcome of an interspecific interaction as a result of environmental change can be explained by effects on 1) the density of the interacting species and/or 2) the traits (including behavior) of the interacting species. These effects in turn can stem directly from altered abiotic conditions (e.g. changes to light or water availability influencing the density or traits of the relevant species), or be mediated by changes in the wider community (i.e., abiotic conditions influence the density or traits of other species in the community, which in turn impinge on the interaction in question) (Fig. 1A; Poisot et al., 2015). In this study, we found that difference in nectar robbing intensity (NRI) of *O. cuspidatum* between a semi-natural and an anthropogenic habitat was primarily the result of partner trait differences, with a lesser contribution of density differences. From the plant’s perspective, these trait differences are driven directly by differing environmental conditions between habitats, while on the robber’s side, trait differences are primarily the result of altered biotic community context (Fig. 1B). Specifically, we found that, together, 1) a nonlinear response of PNRs to the availability of alternative floral resources, 2) greater flower production for *O. cuspidatum* growing in coffee, and 3) higher density of foraging PNRs and greater preference for *O. cuspidatum* by PNRs in the coffee farm than in adjacent forest fragments can fully explain differences in NRI between habitats.

Of these three factors, floral neighborhood composition had the greatest influence on NRI (Table 2). The moderate floral densities found in the coffee fields likely attract more foraging PNRs than found in the low-floral-density forest fragments, leading to higher PNR density and therefore higher NRI for coffee-growing plants (Pathway B in Fig. 1). Since PNR nest density was equivalent between habitats, this indicates preferential foraging by PNRs in coffee fields over forest fragments (the small size of forest fragments in the study area means that bees leaving a nest in the forest can readily move into coffee fields to forage). This finding highlights the importance of shade coffee as a habitat for stingless bees, a result consistent with other studies (Fisher et al., 2017; Jha & Dick, 2010). Moreover, it confirms the importance of floral context as a driver of spatiotemporal heterogeneity in nectar robbery, a relationship which has been hypothesized elsewhere (Irwin & Maloof, 2002), though not explicitly evaluated.

Differences between habitats in *O. cuspidatum* floral traits (flower number and perhaps nectar volume) contributed to higher NRI for plants growing in coffee (Pathway D in Fig. 1). Across most of the observed range of floral display size, more flowers led to more NR. Plants growing in coffee produced more flowers on average [likely because higher light availability increased the amount of photosynthate that plants could allocate to flower production (Fitch & Vandermeer, 2020)], and therefore attracted more PNRs. In addition, PNR foraging behavior differed between habitats, with individual PNR visits lasting longer to flowers in coffee than those in forest. This may be due to differences in nectar rewards between habitats: per-flower nectar volume was 50% higher in coffee than in forest, though this difference was not significant.

Both flower-level and plant-level data indicate that NR significantly reduces fruit set in *O. cuspidatum*. The negative correlation between fruit set and NRI is striking, given that PNR attraction to *O. cuspidatum* is associated with floral traits – i.e. floral display size and nectar quantity – that are frequently positively correlated with both attractiveness to pollinators (Adler & Bronstein, 2004; Theis et al., 2014) and the availability of resources to allocate to reproduction (Bazzaz et al., 1987, 2000). In this population, negative effects of NR are sufficient to outweigh any benefits of increased pollination and/or resource availability, resulting in net reduction in fruit set. However, this effect does not translate to the number of seeds produced per plant, likely because NRI is positively related to flower number; even if a smaller proportion of flowers produces seeds, the net outcome is a consistent number of seeds across a range of NRIs. Thus, despite the difference in NRI intensity between habitats, there was ultimately no difference in reproductive output between plants growing in forest and those growing in coffee (Table 3).

These results suggest that spatial variation in NRI is due primarily to the effects of environmental conditions on the plant community – both on the availability of alternative floral resources and on traits mediating nectar robber attraction. However, whether these results can be generalized to other instances of nectar robbery remains to be seen. More generally, this study highlights the importance of trait differences, rather than or in addition to differences in density, as determinants of interaction frequency or outcome across environmental gradients. This is in line with a large body of research demonstrating that trait-mediated indirect interactions are often more important than density-mediated indirect interactions in determining the effect of one species on another species with which it does not directly interact (Schmitz et al., 2004; Werner & Peacor, 2003). Further, this suggests that the importance of trait-mediated effects of anthropogenic environmental change – on species interactions and, ultimately, species persistence and ecosystem function – have been underappreciated. As this work highlights, detecting and understanding such trait-mediated effects may require fine-grained analysis of interacting organisms across environmental contexts. Thus, on the one hand we heartily endorse recent calls to use network approaches to better understand how environmental change can impact communities beyond just species loss (Poisot et al., 2015, 2017; Tylianakis et al., 2008; Valiente-Banuet et al., 2015). On the other, we caution that the empirical data upon which networks are built often lack the level of detail needed to describe the effects of environmental change on biotic communities.

## Acknowledgements

We are grateful to Walter and Bernd Peters for permission to conduct research at Finca Irlanda. We thank members of the Perfecto and Vandermeer lab groups for support in developing this project. In particular, thanks to Lauren M. Schmitt and Chatura Vaidya for helpful comments on this manuscript, and to Gabriel Dominguez Martinez, Gustavo López Bautista and Chatura Vaidya for support in the field. This study was funded by a Nancy W. Walls Award from the University of Michigan Department of Ecology and Evolutionary Biology and a University of Michigan International Institute Rackham International Research Award, both to Gordon Fitch.

